# *Escherichia coli* O157:H7 responds to phosphate starvation by modifying LPS involved in biofilm formation

**DOI:** 10.1101/536201

**Authors:** Philippe Vogeleer, Antony T. Vincent, Samuel M. Chekabab, Steve J. Charette, Alexey Novikov, Martine Caroff, Francis Beaudry, Mario Jacques, Josée Harel

**Affiliations:** Groupe de Recherche sur les Maladies Infectieuses en Production Animale, Centre de Recherche en Infectiologie Porcine et Avicole, Faculté de médecine vétérinaire, Université de Montréal, Saint-Hyacinthe, QC, Canada; Institut de biologie intégrative et des systèmes, Faculté des sciences et de génie, Université Laval, Québec, QC, Canada; Food and Bioproduct Sciences, College of Agriculture and Bioresources, University of Saskatchewan, Saskatoon, SK, Canada; LPS-BioSciences, Campus d’Orsay, Université Paris-Saclay, Orsay, France; Groupe de Recherche en Pharmacologie Animale du Québec, Département de Biomédecine Vétérinaire, Faculté de Médecine Vétérinaire, Université de Montréal, Saint-Hyacinthe, QC, Canada; Op+Lait, Regroupement de recherche pour un lait de qualité optimale, Faculté de Médecine Vétérinaire, Université de Montréal, Saint-Hyacinthe, QC, Canada

**Keywords:** Keyword: EHEC, biofilm, phosphate, LPS, Pho regulon

## Abstract

In open environments such as water, enterohemorrhagic *Escherichia coli* O157:H7 responds to inorganic phosphate (Pi) starvation by inducing the Pho regulon controlled by PhoB. The phosphate-specific transport (Pst) system is the high-affinity Pi transporter. In the Δ*pst* mutant, PhoB is constitutively activated and regulates the expression of genes from the Pho regulon. In *E. coli* O157:H7, the Δ*pst* mutant, biofilm, and autoagglutination were increased. In the double-deletion mutant Δ*pst* Δ*phoB*, biofilm and autoagglutination were similar to the wild-type strain, suggesting that PhoB is involved. We investigated the relationship between PhoB activation and enhanced biofilm formation by screening a transposon mutant library derived from Δ*pst* mutant for decreased autoagglutination and biofilms mutants. Lipopolysaccharide (LPS) genes involved in the synthesis of the LPS core were identified. Transcriptomic studies indicate the influence of Pi-starvation and *pst* mutation on LPS biosynthetic gene expression. LPS analysis indicated that the O-antigen was deficient in the Δ*pst* mutant. Interestingly, *waaH*, encoding a glycosyltransferase associated with LPS modifications in *E. coli* K-12, was highly expressed in the Δ*pst* mutant of *E. coli* O157:H7. Deletion of *waaH* from the Δ*pst* mutant and from the wild-type strain grown in Pi-starvation conditions decreased the biofilm formation but without affecting LPS. Our findings suggest that LPS core is involved in the autoagglutination and biofilm phenotypes of the Δ*pst* mutant and that WaaH plays a role in biofilm in response to Pi-starvation. This study highlights the importance of Pi-starvation in biofilm formation of E. coli O157:H7, which may affect its transmission and persistence.

**IMPORTANCE:** Enterohemorrhagic *Escherichia coli* O157:H7 is a human pathogen responsible for bloody diarrhea and renal failures. In the environment, O157:H7 can survive for prolonged periods of time under nutrient-deprived conditions. Biofilms are thought to participate in this environmental lifestyle. Previous reports have shown that the availability of extracellular inorganic phosphate (Pi) affected bacterial biofilm formation; however, nothing was known about O157:H7 biofilm formation. Our results show that O157:H7 membrane undergoes modifications upon PhoB activation leading to increased biofilm formation. A mutation in the Pst system results in reduced amount of the smooth type LPS and that this could influence the biofilm composition. This demonstrates how the *E. coli* O157:H7 adapts to Pi starvation increasing its ability to occupy different ecological niches.

## INTRODUCTION

The human enteric pathogen Enterohemorrhagic *E. coli* O157:H7 (EHEC) is responsible for severe collective food-borne infections. Introducing and then consuming contaminated food into production chains contaminates humans. Cattle, the most important reservoir of pathogenic EHEC, are the main agents of its dissemination into the environment (1). Although fully equipped to infect and colonize the host’s intestine, *E. coli* O157:H7 must also be able to respond to changing environmental conditions and to adapt its metabolism to nutrient availability (2).

Phosphorus, one of the most important elements for bacterial growth and survival is 3% of the total dry weight of bacterial cells and the fifth main element after carbon (50%), oxygen (20%), nitrogen (14%), and hydrogen (8%) (3, 4). At the cellular level, phosphorus is in its ionic form called phosphate (PO_4_^3-^) and inorganic phosphate (Pi) is the preferred phosphate source (5). Phosphate is found in many different components of the cell including cell membrane (phospholipids), nucleic acids (the backbone of DNA or RNA), proteins, and complex sugars (e.g. lipopolysaccharides [LPS]). It is also involved in signal transduction (phosphotransfer) and energy production (ATP) (6). Due to its importance for bacteria, phosphate homeostasis is tightly regulated by the two-component regulatory system PhoBR, in which PhoR is a histidine kinase and PhoB is the response regulator controlling expression of genes belonging to the Pho regulon. When the extracellular Pi concentration is high (Pi>4μM), PhoB is inactive and phosphate (Pho) regulon is at the basal expression level, avoiding the use of more complex forms of phosphate. With some environmental changes like nutrient availability and age of culture; and extracellular Pi concentration decreases (below 4 µM) PhoB is activated and regulates the expression of Pho regulon gene leading to changes in phosphate metabolism and transport. Under phosphate-limiting conditions, phosphate transport is controlled by the Pst system encoded by *pstSCAB-phoU* operon belonging to the Pho regulon (6). Any deletion of the *pstSCAB-phoU* operon constitutively activates PhoB leading to constitutive expression of the Pho regulon that results in mimicking phosphate-starvation conditions (7).

In addition to being involved in phosphate homeostasis, the Pho regulon has been connected to bacterial virulence and biofilm formation in several species including *Pseudomonas aeruginosa, P. aureofaciens, P. fluorescens, Agrobacterium tumefaciens, Vibrio cholerae, Bacillus anthracis*, and *Cronobacter sakazakii* (8–15). Recent studies established that *E. coli* O157:H7 strains are able to form biofilms in different environments such as food surfaces, processing plants, and water (16, 17). Although EHEC biofilm formation appears to be strain-dependent, low-temperature conditions favor its formation (18, 19). Stress conditions including nutrient starvation or low oxygen influence biofilm formation by altering the oxidative stress leading to increased release of Shiga-toxin linking pathogenicity, biofilm formation, and stress (20).

In this study, we hypothesized that in EHEC, activation of the regulator PhoB controls the expression of genes that are involved in biofilm formation. Therefore, we investigated biofilm formation of EHEC under phosphate-starvation conditions. By screening a mutant library for decreased ability to agglutinate and to form a biofilm, genes in LPS biosynthetic pathways were over-represented. Further analysis revealed that LPS of Δ*pst* mutant is lacking O-antigen repeats but that its core oligosaccharide was necessary for biofilm formation and agglutination. The glycosyltransferase coding gene *waaH* contributes to Pho regulated biofilm formation. This study highlighted the importance of phosphate-starvation conditions in EHEC biofilm formation that may impact its persistence.

## RESULTS

### Pi limitation promotes biofilm lifestyle of *E. coli* O157:H7

To investigate the effect of Pi starvation on *E. coli* O157:H7 biofilm formation, strain EDL933 was cultivated in high Pi (1.32 mM) or low Pi (1 µM) media in static conditions at 30°C. In low Pi conditions, the relative biofilm formation of EDL933 was 6.15 higher than in high Pi (Fig. 1A). This shows that in low Pi conditions *E. coli* O157:H7 biofilm lifestyle favor being in a biofilm over a planktonic lifestyle like floating as single cells in water (Fig. S1). In previous work, we established that under low Pi conditions, PhoB was activated (21). This allows us to suggest that PhoB could regulate the expression of genes that are related to biofilm formation.

**FIG 1.**
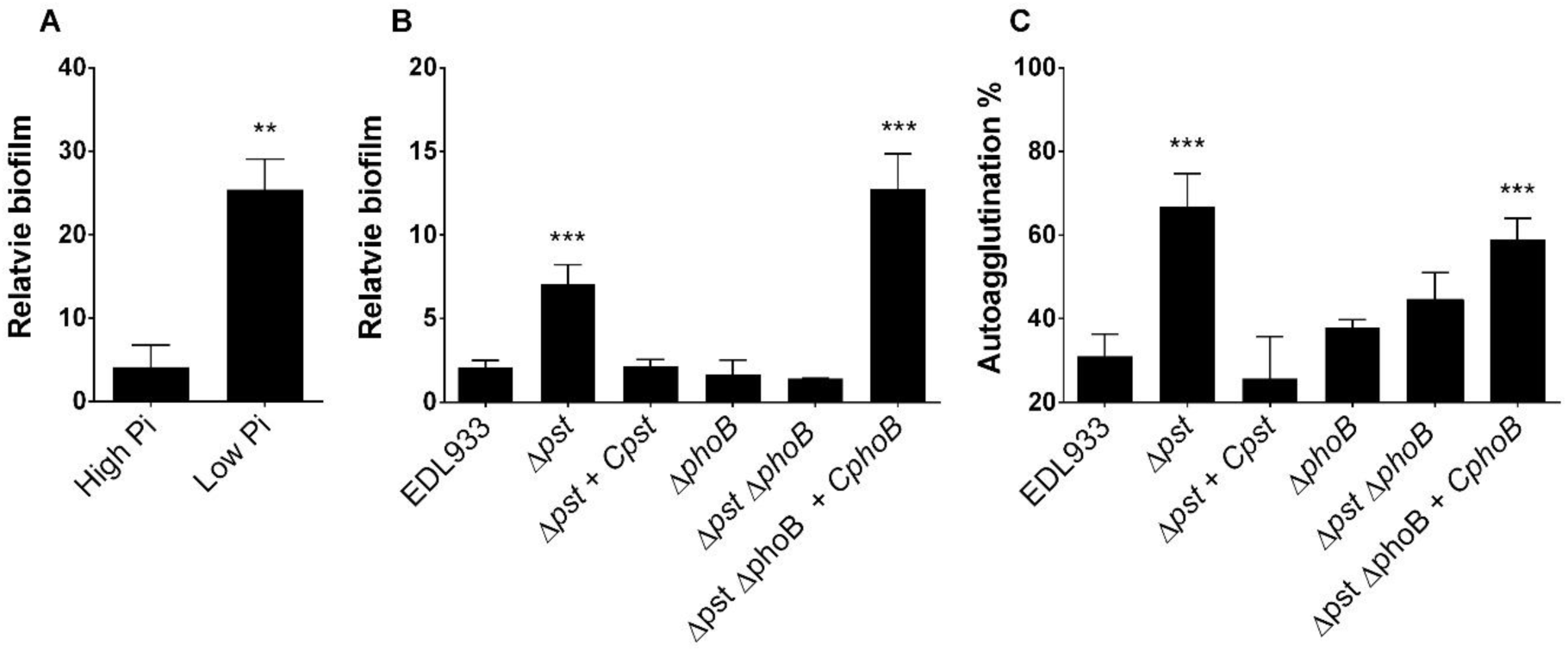
Biofilm and autoagglutination are increased when PhoB is activated. (A) Biofilm of EDL933 formed in MOPS medium containing either 1.32 mM (high Pi) or 1 µM (low Pi) of KH_2_PO. (B) Biofilm formation of EDL933 and its isogenic mutants were formed in M9. The biofilms were evaluated under static conditions in microtiter plates at 30°C and stained with crystal violet. Relative biofilm formation corresponds to biofilm formation (A_595_)/planktonic growth (DO_600_). (C) Autoagglutination was measured after 24 h of growth in M9. Results are averages from at least 3 biological replicates. Statistical analysis was performed using student’s t-test (Fig 1A) or one-way ANOVA with Tukey’s multiple comparison test (Fig 1B and 1C). *\*\* P<0.01; *** P<0.001.*

### Deletion of *pst* increases biofilm formation and autoagglutination

To further investigate the contribution of the Pho regulon in biofilm formation and autoagglutination, we used a series of mutants in which the Pho regulon is constitutively activated (Δ*pst* mutant) or Pho regulon is not activated (Δ*phoB* single mutant and Δ*phoB* Δ*pst* double mutant) (Table 1). The Pho regulon activation status of mutants was confirmed by measuring alkaline phosphatase reporter activity (Fig. S2).

**TABLE 1.**
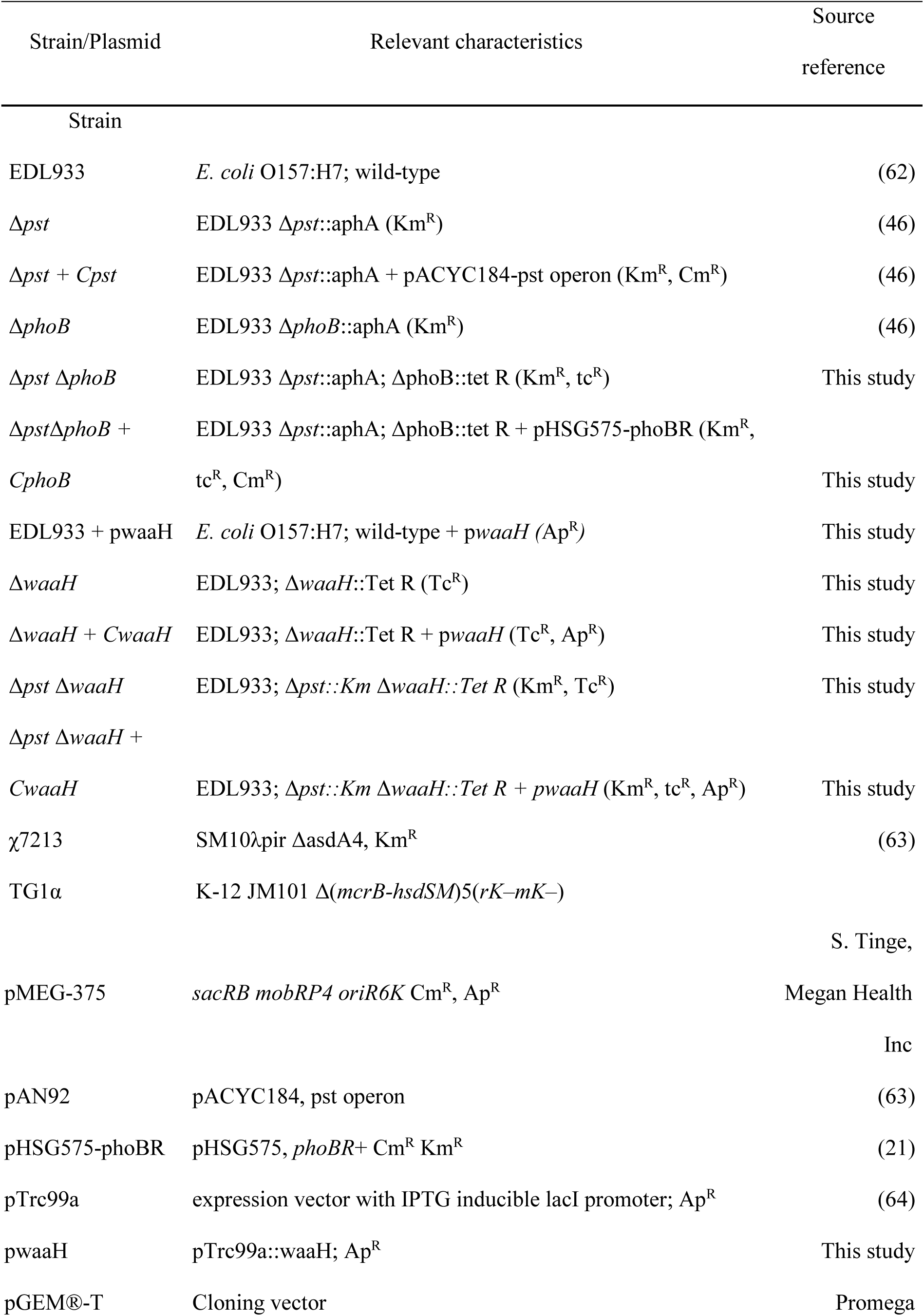
Strains and plasmids used in this study.

| Strain/Plasmid | Relevant characteristics | Source<br>reference |
| --- | --- | --- |
| **Strain** |  |  |
| EDL933 | *E. coli* O157:H7; wild-type | (62) |
| $\Delta pst$ | EDL933 $\Delta pst::aphA$ (Km<sup>R</sup>) | (46) |
| $\Delta pst + Cpst$ | EDL933 $\Delta pst::aphA$ + pACYC184-*pst* operon (Km<sup>R</sup>, Cm<sup>R</sup>) | (46) |
| $\Delta phoB$ | EDL933 $\Delta phoB::aphA$ (Km<sup>R</sup>) | (46) |
| $\Delta pst \Delta phoB$ | EDL933 $\Delta pst::aphA$; $\Delta phoB::tet R$ (Km<sup>R</sup>, tc<sup>R</sup>) | This study |
| $\Delta pst \Delta phoB +$<br>*CphoB* | EDL933 $\Delta pst::aphA$; $\Delta phoB::tet R$ + pHSG575-*phoBR* (Km<sup>R</sup>,<br>tc<sup>R</sup>, Cm<sup>R</sup>) | This study |
| EDL933 + *pwaaH* | *E. coli* O157:H7; wild-type + *pwaaH* (Ap<sup>R</sup>) | This study |
| $\Delta waaH$ | EDL933; $\Delta waaH::Tet R$ (Tc<sup>R</sup>) | This study |
| $\Delta waaH + CwaaH$ | EDL933; $\Delta waaH::Tet R$ + *pwaaH* (Tc<sup>R</sup>, Ap<sup>R</sup>) | This study |
| $\Delta pst \Delta waaH$ | EDL933; $\Delta pst::Km \Delta waaH::Tet R$ (Km<sup>R</sup>, Tc<sup>R</sup>) | This study |
| $\Delta pst \Delta waaH +$<br>*CwaaH* | EDL933; $\Delta pst::Km \Delta waaH::Tet R$ + *pwaaH* (Km<sup>R</sup>, tc<sup>R</sup>, Ap<sup>R</sup>) | This study |
| $\chi 7213$ | SM10λpir $\Delta asdA4$, Km<sup>R</sup> | (63) |
| TG1α | K-12 JM101 $\Delta(mcrB-hsdSM)5(rK-mK-)$ | |
| **Plasmid** |  |  |
|  |  | S. Tinge, |
| pMEG-375 | *sacRB mobRP4 oriR6K* Cm<sup>R</sup>, Ap<sup>R</sup> | Megan Health |
|  |  | Inc |
| pAN92 | pACYC184, *pst* operon | (63) |
| pHSG575-phoBR | pHSG575, *phoBR*+ Cm<sup>R</sup> Km<sup>R</sup> | (21) |
| pTrc99a | expression vector with IPTG inducible *lacI* promoter; Ap<sup>R</sup> | (64) |
| pwaaH | pTrc99a::*waaH*; Ap<sup>R</sup> | This study |
| pGEM®-T | Cloning vector | Promega |

The relative biofilm formation and the autoagglutination phenotype of Δ*pst* mutant in M9 conditions were significantly increased compared to EDL933 (Fig. 1B and 1C). Low biofilm level was restored to wild-type status in a *pst-*complemented strain. Single deletion of *phoB* appeared to have no impact on either biofilm formation or on autoagglutination. However, the double-deletion mutant, Δ*phoB* Δ*pst*, demonstrated low ability to form a biofilm (Fig. 1B). These results correlated with the Pho regulon activation (Fig. S2). Consequently, the activation of PhoB is required to enhanced agglutination and biofilm formation of a Δ*pst* mutant.

### The Δ*pst* mutant biofilm contains N-acetylglucosamine (GlcNAc)

The higher biofilm formation of the Δ*pst* mutant was observed by confocal laser scanning microscopy (CLSM) with the FM1-43 stain confirming the crystal violet data (Fig. 2A). Moreover, while Δ*pst* mutant appeared to form a much denser biofilm compared to EDL933, *pst*-complemented strains and double-mutant Δ*pst* Δ*phoB*, no difference in thickness was observed. (Fig. 2A). Also, while growth in a planktonic form was almost abolished in low Pi conditions (Fig. S1B), biofilm density and thickness were similar to high Pi conditions (Fig. 2B). This last point highlighted that in low Pi conditions, biofilm lifestyle is favored over planktonic lifestyle. By using fluorophores that stained proteins (Sypro), cellulose (Calcofluor), or N-acetylglucosamine (GlcNAc) residues (WGA), biofilm was characterized. Protein and cellulose were not detected in any biofilm matrices (data not shown). Interestingly, GlcNAc residues were detected in biofilm matrix formed by Δ*pst* mutant and absent in those formed by EDL933 and Δ*pst* + *pst*-complemented strain. Notably, as GlcNAc residues were also detected in Δ*pst ΔphoB* double-mutant matrix, the indication is that this characteristic of the matrix is not due to PhoB activation. The absence of GlcNAc in the biofilm matrix of wild type grown in low Pi conditions (Fig. 2B) is confirmed. Taken together, CLSM results confirmed that Δ*pst* mutant formed denser biofilm than wild type and that GlcNAc residues are contained in its matrix.

**FIG 2.**
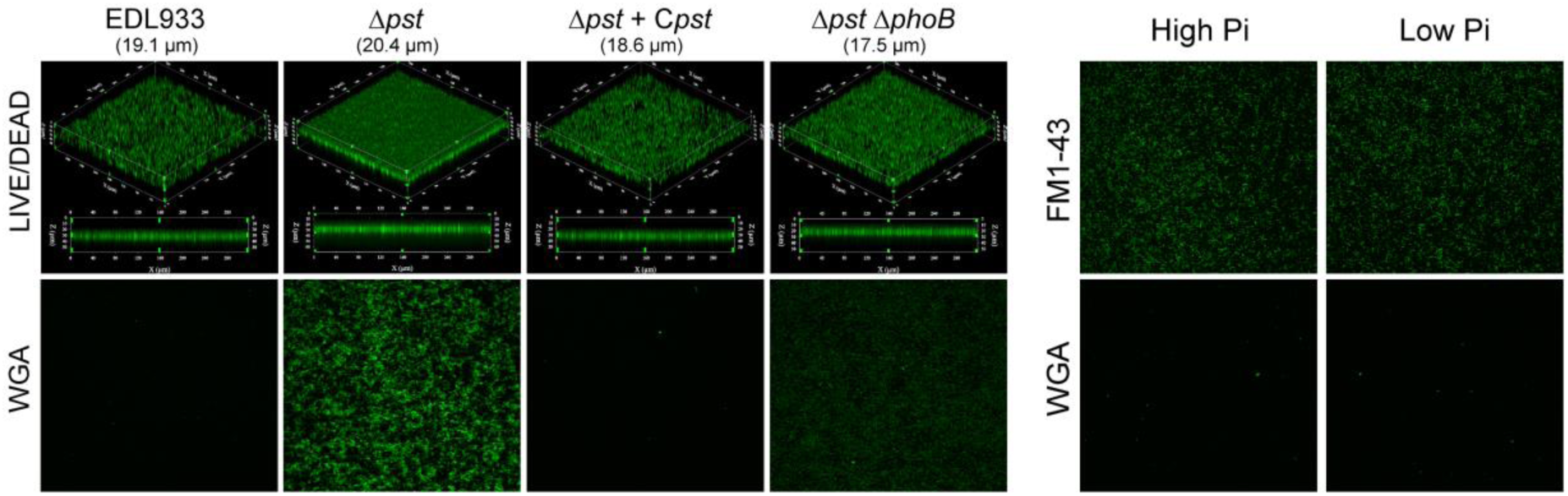
Δ*pst* mutant forms denser biofilm than wild type and exposes GlcNAC residues. Confocal laser scanning microscopy images of 24-h biofilms formed by EDL933 and its derivative mutants. (A) Projections of Z-stacks of biofilm stained with FM1-43 and photographs of biofilm stained with WGA. (B) Photos of EDL933 biofilms grown in MOPS high or low Pi, and stained with FM1-43 or WGA.

### Identification of putative Pho regulon members involved biofilm formation

To identify Pho regulon members involved in enhanced biofilm and agglutination phenotypes of Δ*pst* mutant, a mini-Tn*10* transposon mutant library was generated from the Δ*pst* mutant. A total of 5118 mini-Tn*10* Cm^R^ mutants were screened for their reduced ability to autoagglutinate. From a total of 122 mutants displaying an autoagglutination phenotype different from Δ*pst* mutant, 93 (1.8% of the library) showed reduced ability to form a biofilm. Using high-throughput sequencing, 88 transposon insertion sites were identified (Table S2). Targeted genes were regrouped by categories of orthologous groups (COG) (Fig. 3). While the majority of genes had unknown functions, a high proportion of mutations associated with decreased biofilm formation was located in genes affecting the membrane and cell wall biosynthesis including genes of the LPS R3 core synthesis (*waa* locus).

**FIG 3.**
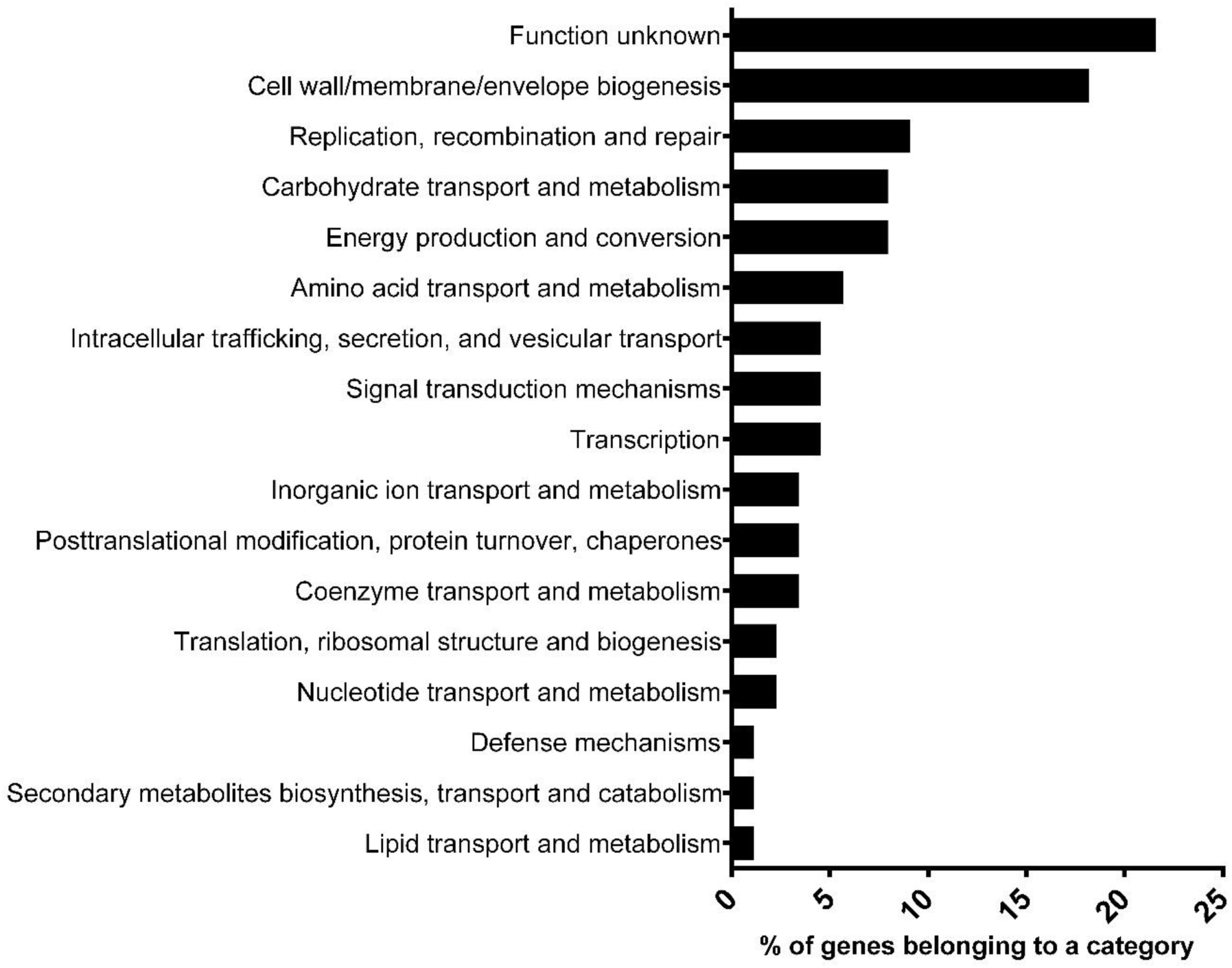
Biofilm and autoagglutination transposon mutants are mainly involved in cell wall synthesis. COG analysis with eggNOG-mapper revealed that transposons of autoagglutination and biofilm negative mutants were inserted mainly in genes coding for cell wall synthesis and unknown function.

### Transposon insertion sites of several mutants are in LPS core synthesis genes

Five independent insertions that resulted in the loss of biofilm formation were identified in *waaC, waaE, waaF, waaG*, and *waaQ* from the LPS core oligosaccharide biosynthesis (*waa*) locus (Fig. 4A–C). In *E. coli* O157, the majority of enzymes required for core oligosaccharide assembly are encoded by genes within the chromosomal *waa* locus (22). The mutation sites were in genes *waaE (*also named *hldE*), involved in the synthesis of the heptose precursor (23); in genes *waaF* and *waaC* (encoding by the *rfaD-waaFCL* operon) that are responsible for the addition of heptose I and II on the LPS core; and in genes *waaQ* and *waaG (*encoding by the *waaQGP* operon) involved in the addition of Heptose III and glucose I on the LPS core (Fig. 4A and 4B and Table S2). This result strongly suggests that LPS core synthesis is important for autoagglutination and biofilm formation of the Δ*pst* mutant. Moreover, the biofilm matrices of those Tn*10* mutants were negative to WGA staining (Fig. 4D), suggesting that an intact inner and outer LPS core structure is necessary for biofilm formation and that the detection of GlcNAc on the biofilm matrix is linked to the LPS structure of Δ*pst* mutant.

**FIG 4.**
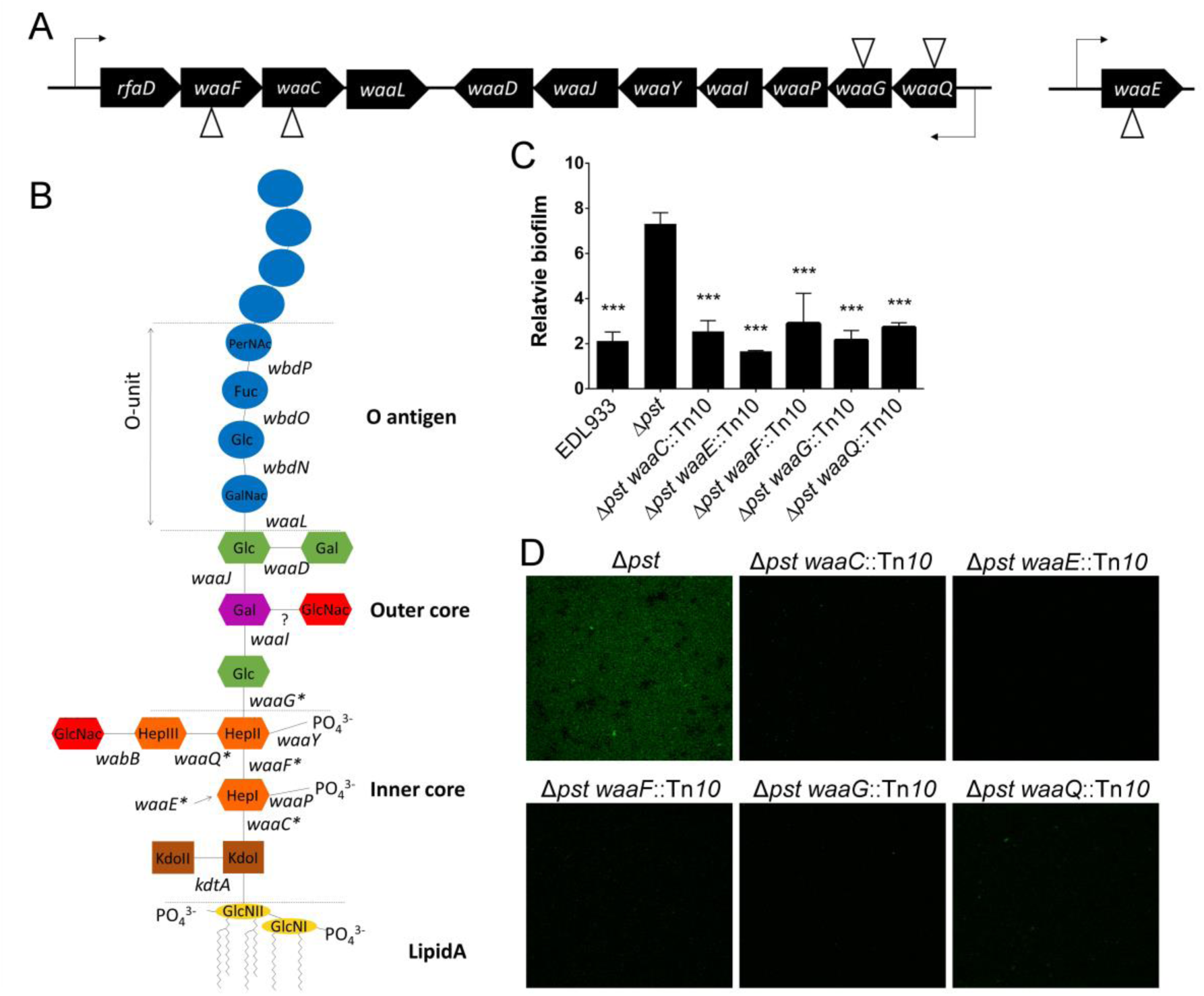
O157 LPS synthesis is important for biofilm formation of Δ*pst* mutant. (A) Genetic organization of LPS core biosynthesis locus. Tn*10* insertions sites are indicated by triangles (B). Structure of O157 LPS structure. Genes with stars show genes where a transposon was inserted. Gene *waaE* is involved in the synthesis of ADP-L-glycero-β-D-manno-heptose, the heptose substrate for *waaC, waaF*, and *waaQ*. (C) Relative biofilm formation of Δ*pst*-derivative mutants with Tn*10* inserted in LPS synthesis genes in M9 at 30°C for 24 h. Relative biofilm formation corresponds to biofilm formation (A_595_)/planktonic growth (DO_600_). Results are averages from at least 3 biological replicates. Statistical analysis was performed using a one-way ANOVA with Tukey’s multiple comparison post-test. Stars indicate significant results compared to Δ*pst* mutant *** *P* < 0.001. (D) Confocal laser scanning microscopy images of 24-h biofilms stained with WGA of EDL933, Δ*pst* mutant, and Δ*pst* derivative mutant with Tn*10* inserted in LPS synthesis genes grown in M9 at 30°C for 24 h.

Concomitantly our transcriptomic studies supported the influence of *pst* mutation and Pi starvation on the LPS biosynthetic pathway. Many of its genes involved in lipid A, core and O-unit biosynthesis and export were downregulated in both the Δ*pst* mutant and Pi starvation conditions (Table 2), suggesting that LPS synthesis could also be under the direct or indirect control of PhoB. *In silico* analysis identified putative Pho boxes in proximity to genes and/or operons involved in the different steps of LPS biosynthesis (Table 2) suggests that genes involved in LPS synthesis could also be under the direct or indirect control of PhoB.

**TABLE 2.**
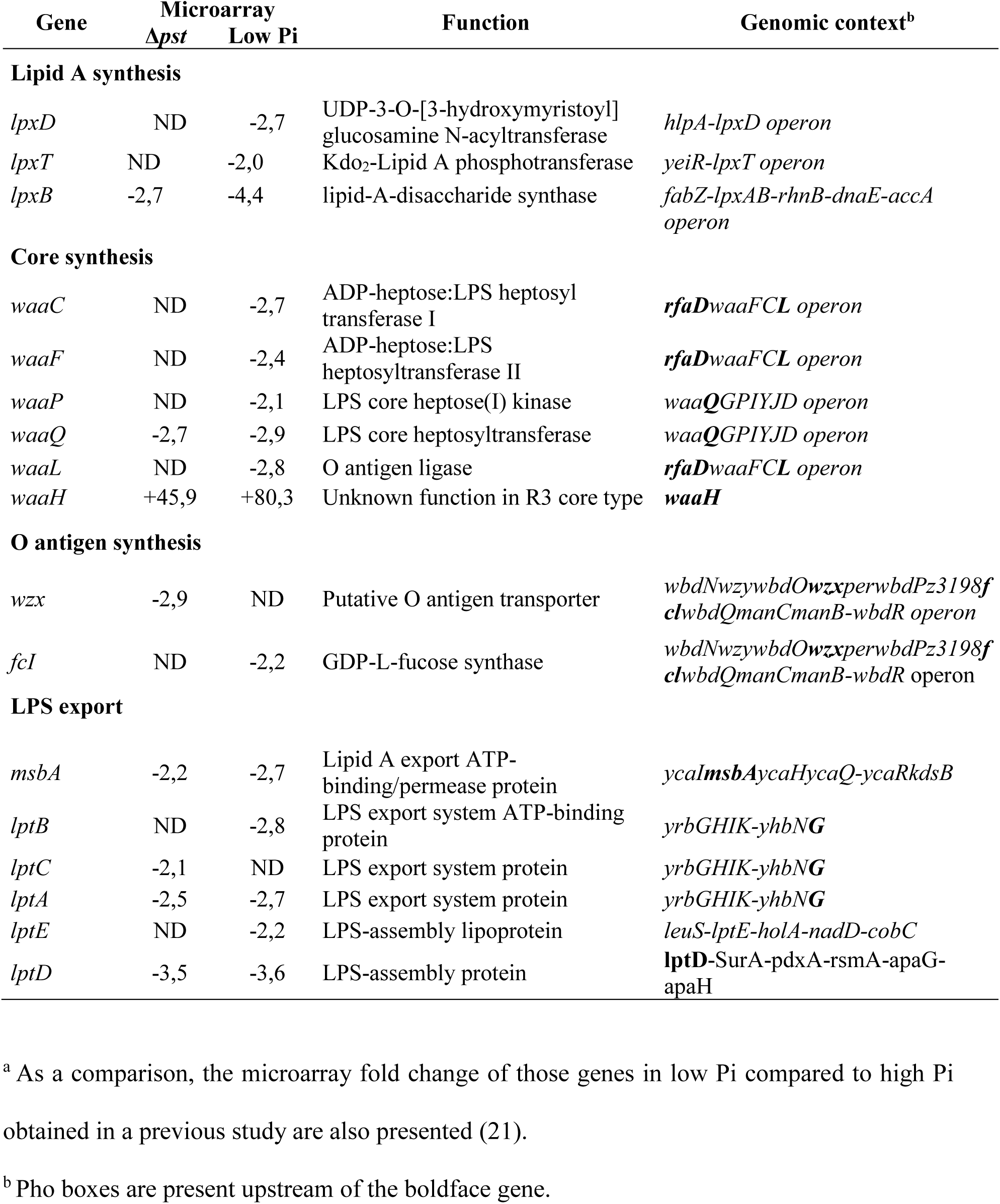
Fold change of genes related to LPS synthesis in the Δ*pst* mutant compared to EDL933.

### O-antigens are absent from Δ*pst* mutant LPS

The LPS pattern of planktonic and biofilm cells of the Δ*pst* mutant were analyzed by SDS-PAGE. The LPS molecule of *E. coli* O157:H7 consists in a R3 type core oligosaccharide (core OS) linked to the hydrophobic lipid A molecule, and being substituted by the long chain O157 antigenic polysaccharide (22). The SDS-PAGE LPS profile was examined to determine whether the *pst* mutation is accompanied by major changes to the LPS structure. LPS preparations were isolated either from cells grown in LB, or from biofilm, and planktonic cells grown in M9. Strain EDL933 and its Δ*pst* mutant differed mainly in the absence of O-specific chains in the mutant. Some differneces were also observed for the stained molecular species migrating to the lower region of the gel correspond to signals in the lower mass region of the MALDI mass spectrum. They are short chain LPS for EDL933 consisting of the lipid A and the core OS, together with the same plus one, two, or three additional repeats of O-units. The electrophoretic profiles of this LPS lower mass region was different in the Δ*pst* mutant. In this case, the core plus lipid A region is present and similar to that observed for the EDL933 strain, i.e. represented by a double band. At the same time, the bands corresponding to additional O-units are absent, instead, a weak additional band is present, migrating slightly less than the lipid A plus core (Fig. 5A). In the EDL933 LB preparation, the upper signal of the double band is very weak. These structural features are accessible to MALDI-MS analysis and will be discussed in details in the MALDI-MS section.

**FIG 5.**
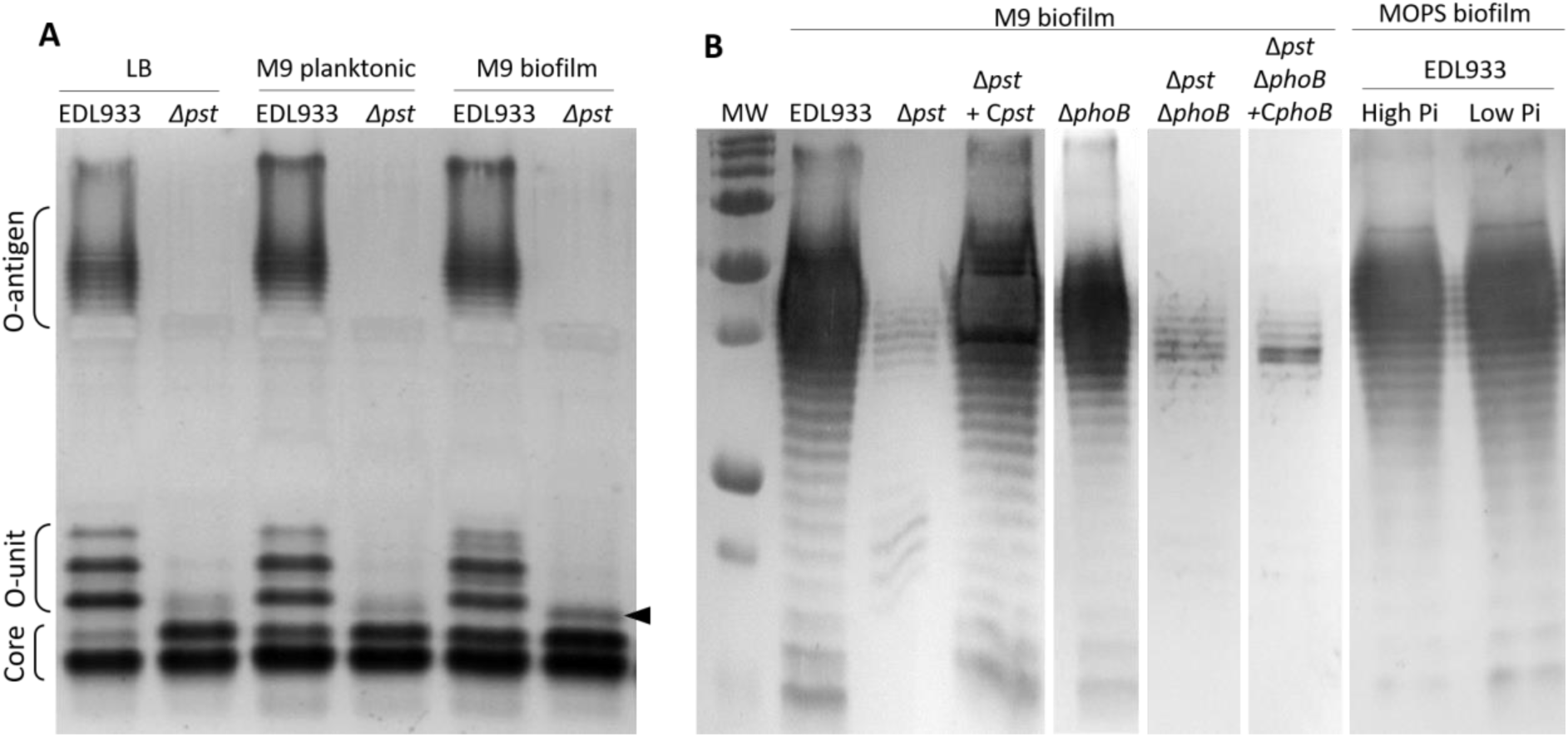
Deletion of *pst* leads to the absence of O157 antigens. (A) LPS of EDL933 and Δ*pst* mutant grown in LB and as planktonic or biofilm cells in M9 at 30°C were extracted, separated on 15% SDS-PAGE and then stained by silver stain. The black arrow shows the presence of an additional slower migrating band in Δ*pst* LPS extract that was absent in EDL933 LPS extract. (B) Western blot analysis of LPS extracts of EDL933 and its derivative mutants grown as biofilm cells in M9 at 30°C or of EDL933 grown as biofilm cells in MOPS high Pi or low Pi. LPS were revealed by using primary O157:H7 monoclonal antibody (1:1000) and a horseradish peroxidase-conjugated antibody (1:1000).

Using Western blot analysis with anti-O157 monoclonal antibodies, O157 LPS specific regular banding pattern was observed in EDL933, however band intensity was very weak in the Δ*pst* mutant (Fig. 5B). Thus, the Δ*pst* mutant predominantly produces rough type LPS species.

To further investigate LPS structure modifications, we used MALDI-TOF mass spectrometry to analyze the LPS extracts of EDL933 and Δ*pst* mutant grown in LB conditions; and from biofilm and planktonic grown in M9 conditions (Fig. 6). We first analyzed the LPS polysaccharide or oligosaccharide moieties, isolated by mild acid hydrolysis. The studied *m/z* range corresponds to the low molecular weight species that were observed in the low mass-region of the SDS-PAGE (Fig. 5A). The negative-ion MALDI mass-spectra of the PS moieties of LPS isolated from different culture conditions of the EDL933 and those from the Δ*pst* mutant cultures are presented in Figure 6. For better clarity, we set up the upper limit of the presented *m/z* range at 2800u, which corresponds to the core plus one repeating O-chain unit. However, molecular ions containing up to three O-chain units were clearly detected for the wild-type strain. In the *m/z* region 1700-2100 of all the spectra presented in Figures 6, peaks corresponding to [M-H]^−^ molecular ions of the core oligosaccharide were observed and will be further detailed. In the 2500-2800u region of all spectra of the EDL933 strain, the same pattern of peaks was observed shifted at 699 mass units. This shift corresponds to the mass of one repeating O-chain units of *E. coli* O157, consisting of -GalNAc-PerNAc-Fuc-Glc-(24). As mentioned above, and in accordance to the SDS-PAGE data, we observed for the wild-type strain, molecular ion peaks corresponding to the addition of one, two and three repeating O-chain units (data not shown). The lack of an O-chain in the Δ*pst* mutant was confirmed by the absence of peaks corresponding to one or two O units (+ 699 u). Instead, for each culture condition, we observed a group of peaks shifted at plus 365 mass units relatively to the peaks of the core oligosaccharide molecular species. This group of peaks could correspond to the weak additional band observed in SDS PAGE for the mutant LPS. The mass of 365u of this additional structural element could be interpreted in different ways, e.g. a disaccharide unit containing a hexose (Hex) plus N-acetyl-hexose (HexNAc) or Hex plus pyrophosphoryl-ethanolamine (PPEA).

**FIG 6.**
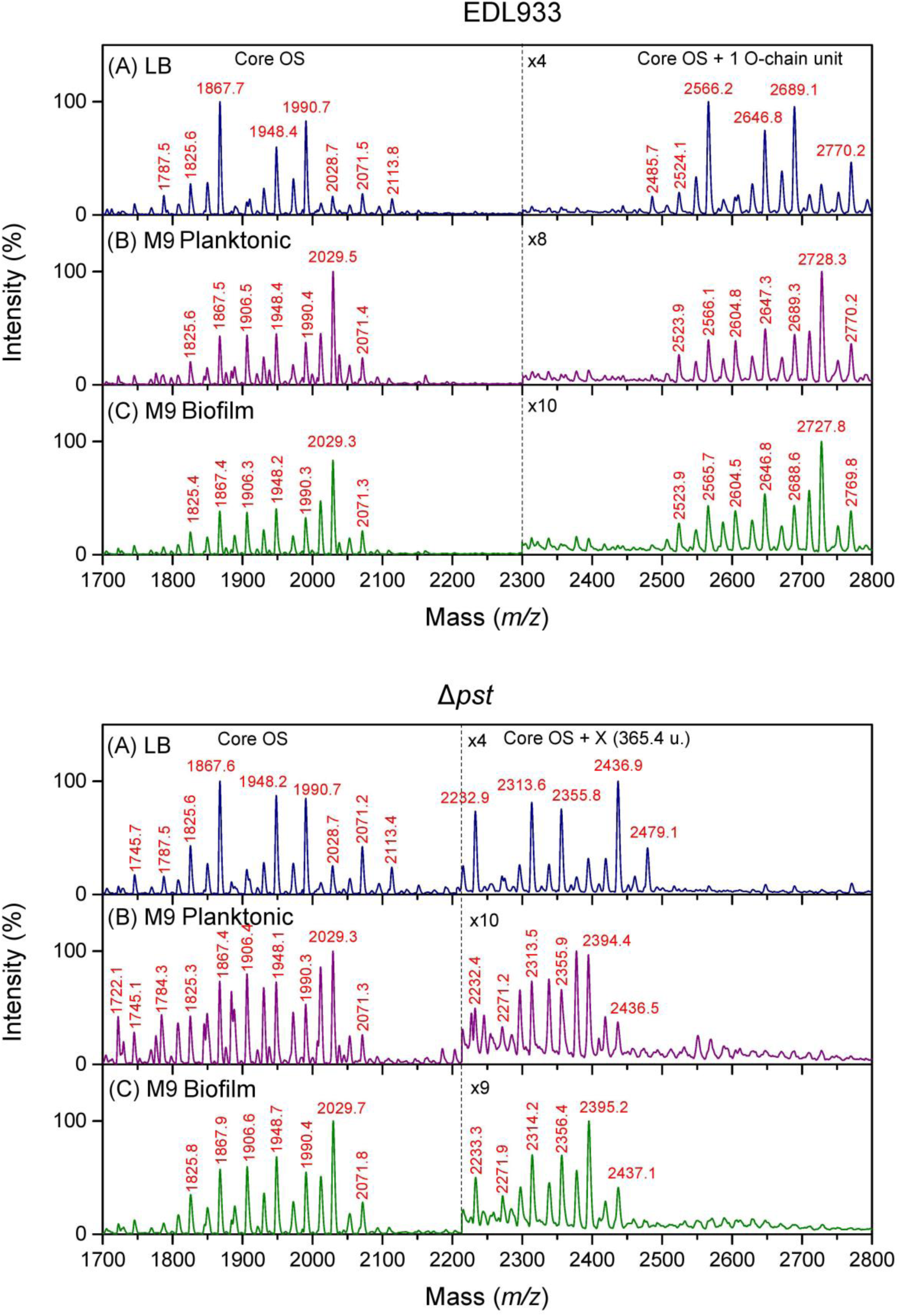
LPS of Δ*pst* mutant are rough LPS lacking O-antigen. Mass spectroscopy analysis of LPS extracted from EDL933 (upper panel) and Δ*pst* mutant (bottom panel) grown in (A) LB or in M9 as planktonic (B) (M9 planktonic) or biofilm (C) (M9 biofilm) reveals the absence of the O chain from LPS of Δ*pst* mutant and presence of an additional peaks shifted at plus 365 mass units from the core peaks, and corresponding to a putative disaccharide. Moreover, this analysis suggests an increase of non-acetylated GlcN in the cores of both strains under M9 culture conditions. To better visualized the mass region between 2200 m/z and 2800 m/z for EDL933 and 2300 m/z and 2800 m/z for Δ*pst* mutant were zoomed-in. Core OS: core oligosaccharide; X: unknown structural element.

### Core oligosaccharide structures varied as a function of culture conditions in a similar way for both strains

LPS core structures corresponded to R3 core type and were found in both EDL933 and Δ*pst* mutant LPS. On the basis of the characterized core structure (22), after hydrolysis, a major peak at 1868 m/z present in all six spectra (Figure 6) corresponds to the core region including a structure with 1 molecule of Kdo (the second Kdo is released during hydrolysis); 3 heptose (Hep); 4 Hex; and 2 HexNAc. For both strains in LB conditions, the major peaks appear at *m/z* 1868, 1948 and 1991. According to Kaniuk and collaborators, the last two peaks correspond respectively to addition of one phosphate (P) or one phosphoryl-ethanolamine (PEA) (22). A minor peak at *m/z* 1825 can be interpreted by the absence of an Ac group on one of the branched GlcN. All these peaks are also observed in spectra corresponding to the M9 cultures of both strains, however they are relatively small. Under these conditions two peaks at *m/z* 1906 and 2029 become major ones for both strains and can be interpreted by addition of P and PEA to the molecular species at *m/z* 1825 lacking one Ac. Such variability has been already reported (22).

### Lipid A structures are different between WT and mutant strains, and further modified according to culture conditions

The negative-ion mass-spectra of lipid A isolated from the EDL933 wild-type strain and the Δ*pst* mutant are presented in Figure 7A. They correspond to the WT and mutant grown under three different culture conditions, i.e. LB culture, M9 planktonic culture, and M9 biofilm culture. Peaks of three major molecular species were observed at *m/z:* 1797.4, 1920.5 and 2035.8, corresponding respectively to the non-modified *E. coli* classical hexa-acyl molecular species, the hexa-acyl molecular species with one phosphate group being substituted with a PEA residue, and a hepta-acyl molecular species with palmitoylation at the secondary C-2 position. The structures of this well-known molecular species are presented in Figure 7B. While the non-modified hexa-acyl molecular species is equally observed in all the six mass spectra, the modified ones are differentially represented according to the different strains, and under different culture conditions.

**FIG 7.**
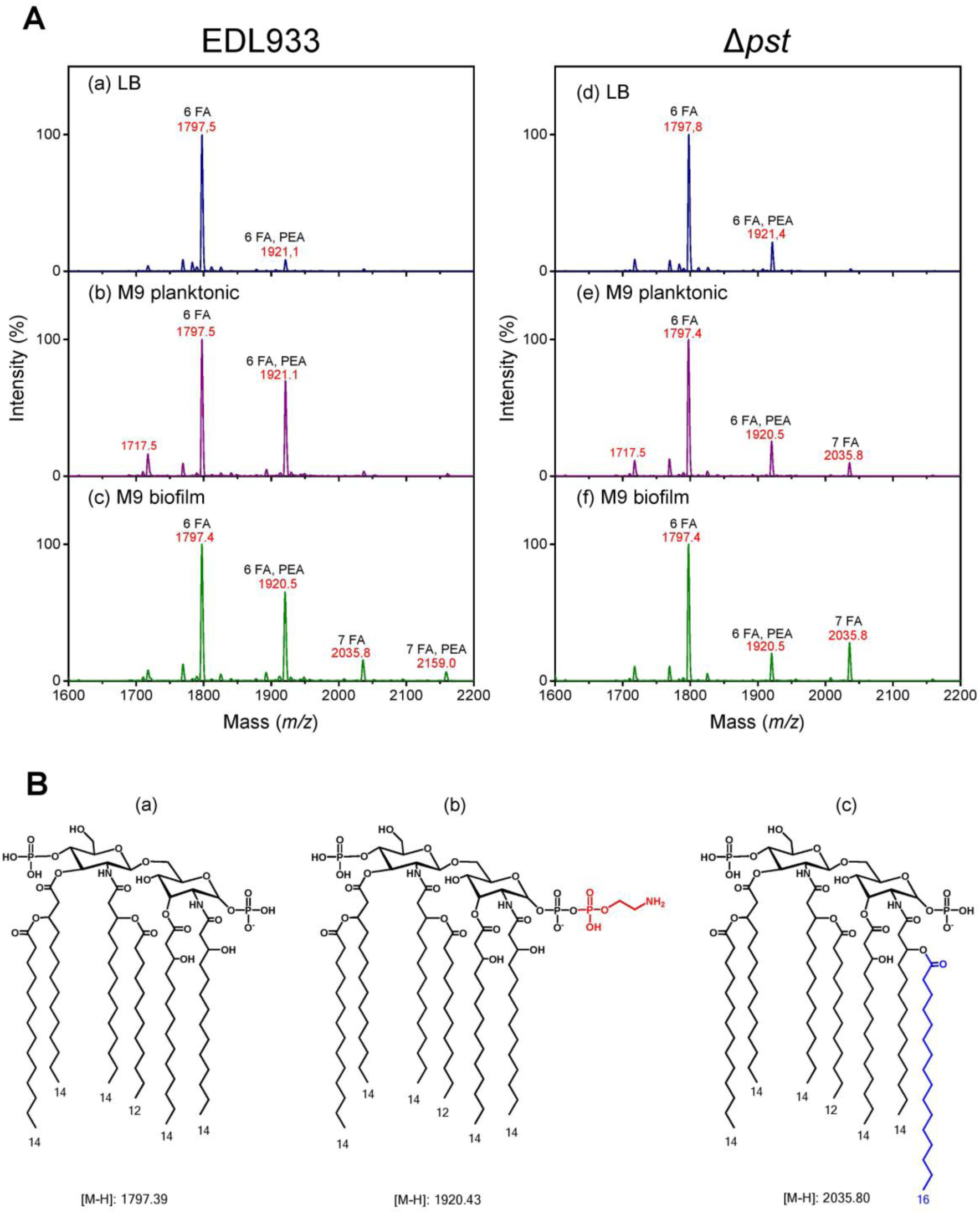
Lipid A structures are different between EDL933 and Δ*pst* mutant, and differentially modified according to culture conditions. (A) Mass spectroscopy analysis of Lipid A from EDL933 (left panel) and Δ*pst* mutant (right panel) grown in (a and d) LB or in M9 as planktonic (b and e) (M9 planktonic) or biofilm (c and f) (M9 biofilm) reveals that PEA substitution is culture dependent for EDL933 and non-dependent for the Δ*pst* mutant. Moreover, palmitoylation is increased under biofilm conditions for both strains. (B) Structure representation of three major molecular species of lipid A corresponding to the hexa-acyl molecular species (A), the hexa-acyl molecular species with one phosphate group being substituted with a PEA residue (B), and an hepta-acyl molecular species with palmitoylation at the secondary C-2 position (C). FA, fatty acid; PEA, phosphorylethanolamine.

Our data suggest that the substitution of the phosphate group with PEA is strongly increased when the wild-type strain is grown in the minimal medium (M9) comparatively to the reach medium (LB). This is not the case for the Δ*pst* mutant for which the presence of the PEA substituted molecular species seems to be independent from growth conditions. It is also somewhat intermediate between the two extreme levels observed for the EDL933 strain, i.e. higher than in the EDL933 LB culture and lower in both EDL933 M9 cultures.

As for the palmitoylated molecular species, it is almost absent in EDL933 LB and M9 planktonic cultures, as well as in the Δ*pst* mutant LB cultures. For the EDL933 strain it is small but fairly detectable under M9 biofilm conditions. For the Δ*pst* mutant, it is small under M9 planktonic conditions, and its presence is considerably increased under M9 biofilm conditions. These observations suggest that lipid A palmitoylation in both strains is favored by the biofilm growth conditions. This is coherent with our earlier findings (25). For the Δ*pst* mutant, the lipid A palmitoylation might be somewhat higher than for the wild-type strain in minimal medium (M9), since it is observed even in the M9 planktonic culture, while it is strongly inhibited in the rich medium (LB) for both strains.

### The regulator PhoB is not involved in the absence of O-antigen

To assess the role of the regulator PhoB in the absence of O-antigen production in the Δ*pst* mutant, LPS from Δ*pst* Δ*phoB* double mutants; and from EDL933 cells grown in low Pi conditions were analyzed by Western blot (Fig. 5B). While the Δ*pst* Δ*phoB* double mutant was deficient in O-antigen, EDL933 made an O-chain antigen when grown in low Pi conditions. Moreover, PhoB complementation of the Δ*pst* Δ*phoB* double mutant did not restore O-antigen (Fig. 5B). These results suggest that the absence of O-antigen is not under PhoB control and is specific to the *pst* mutation. (Fig 1A and 5B).

### *waaH* encoding a glycosyltransferase contributes to biofilm formation

While most of the genes involved in the biosynthesis and export of LPS were all downregulated, the gene *waaH* encoding for a predicted glycosyltransferase was strikingly upregulated in the Δ*pst* mutant (+45.9) and in low Pi conditions (+80.3) (Table 2). Klein *et al*. showed that *waaH* was responsible of the addition of glucuronic acid (GlcUA) on the third heptose of LPS R2 type core in *E. coli* K12 strain but not for the R3 type core of O157 LPS (26–28) (Fig. 6). No modification corresponding to the GlcUA mass (176 u) was observed in Δ*pst* mutant by MALDI-TOF analysis (Fig. 6). No difference in LPS patterns between Δ*waaH* single and Δ*pst* Δ*waaH* double mutants was observed by SDS-PAGE and Western blot (Fig. 8A and Fig. S7). Interestingly, biofilm formation was significantly reduced in the Δ*pst* Δ*waaH* double mutant (Fig. 8B), and in the Δ*waaH* single mutant in low Pi conditions (Fig. 8C). However, the deletion of *waaH* did not affect the autoagglutination phenotype (data not shown). Thus, WaaH plays a role in biofilm formation in a Pho-dependant manner. Pho box consensus sequences were identified *in silico* in the promoter region of *waaH* of *E. coli* O157:H7 EDL933; and EMSA results showed PhoB dose-dependent shifts of the probes designed from the promoter region of *waaH* of EDL933 (Fig. S6A). Finally, Pho-regulated WaaH plays a role in *E. coli* O157:H7 biofilm formation in response to Pi-starvation and in the Δ*pst* mutant and is not associated with GlcUA addition on O157 LPS.

**FIG 8.**
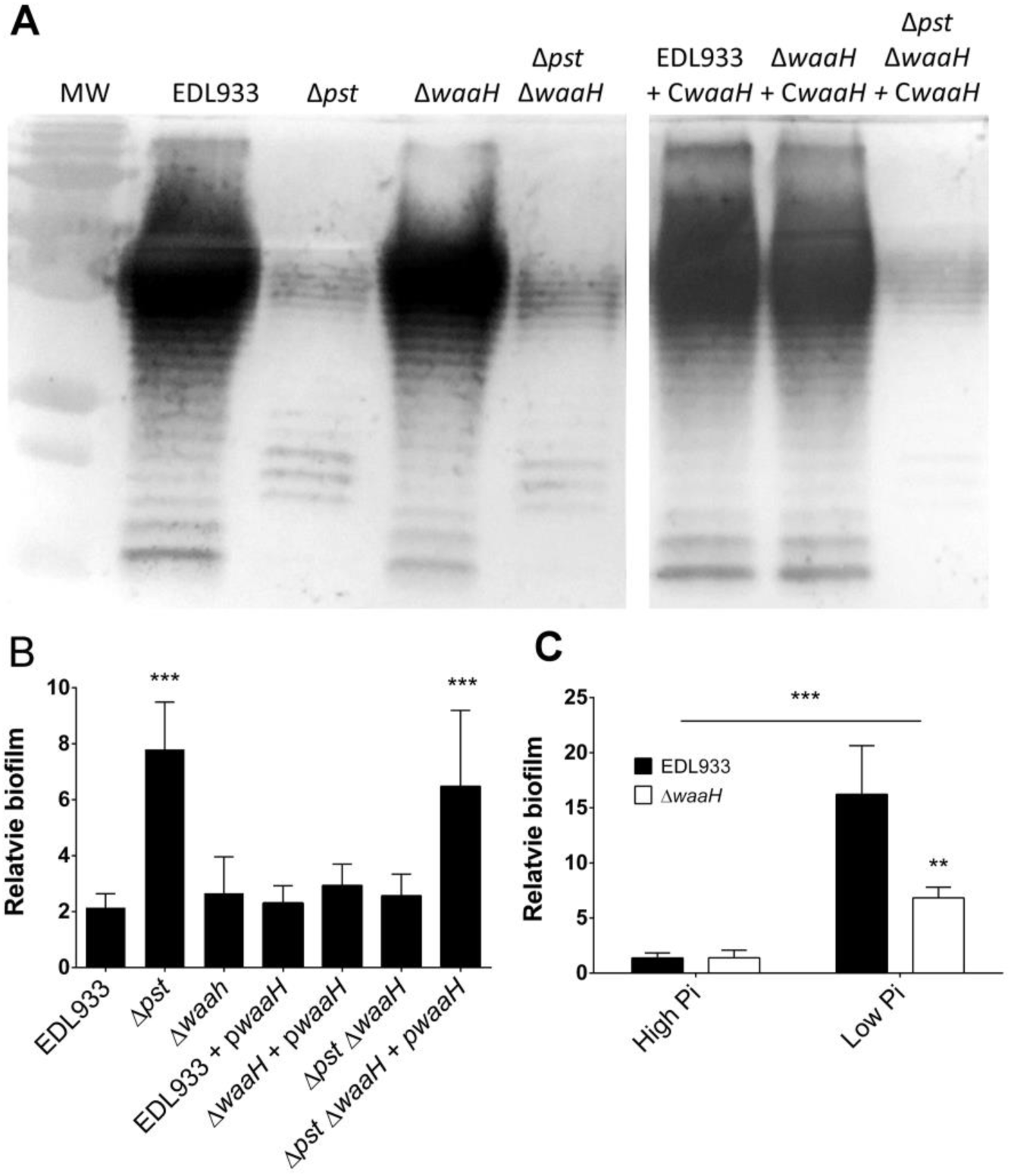
*waaH* played a role in biofilm formation in response to Pi starvation but its function remains unknown. (A) Western blot of LPS extracts of EDL933 and its derivative mutants grown as biofilm cells in M9 at 30°C. (B) Relative biofilm formation of EDL933 and its derivative mutants cultured in M9 at 30°C for 24 h. Results are averages from at least 3 biological replicates. Relative biofilm formation corresponds to biofilm formation (A_595_)/planktonic growth (DO_600_). Statistical analysis was performed using a one-way ANOVA with Tukey’s multiple comparison post-test. Stars indicate significant results compared to Δ*pst* mutant *** *P* < 0.001. (C) Relative biofilm formation of EDL933 and Δ*waaH* grown in MOPS high Pi and low Pi at 30°C for 24 h. Results are averages from at least 3 biological replicates. Statistical analysis was performed using a two-way ANOVA with Bonferroni’s multiple comparison post-test. ** *P* = 0.0012.

## DISCUSSION

### PhoB favors biofilm formation

In this study, we show that activation of the transcriptional regulator PhoB promotes biofilm formation in phosphate-starvation conditions and in the *pst* mutant where PhoB is derepressed. Activation of PhoB has been shown to affect biofilm formation in different bacterial species (8, 9, 11, 12, 14, 15, 29). Upon depletion of phosphate, the plant pathogen *A. tumefaciens* was shown to increase its biofilm formation. This was linked to the higher production of the unipolar polysaccharide adhesion (30). In pathogenic *E. coli*, phosphate limitation was shown to affect adhesion of enteropathogenic *E. coli* (31) and to increase virulence (21). In view of these results, we suggest that PhoB activation through phosphate starvation could have positive or negative effect on biofilm formation depending on species or even pathotype.

### LPS are important for biofilm formation

Our study emphasizes the importance of LPS in biofilm formation and autoagglutination phenotypes of the Δ*pst* mutant as revealed by insertion mutants in genes involved in the core LPS biosynthesis. LPS are surface phosphorylated lipoglycans present in the outer leaflet of the outer membrane of Gram-negative bacteria (32). The LPS of *E. coli* O157:H7 is a tripartite molecule consisting of the hydrophobic anchor lipid A, the core OS, which is divided into the inner and outer core regions, and the O157-antigen polysaccharide (32, 33).

Within the collection of biofilm autoagglutination Tn*10* mutants generated from Δ*pst* mutant, the transposon was inserted in genes within the *waa* locus that are responsible for the enzymes needed for the stepwise assembly of the LPS inner and outer core oligosaccharides (32) (Fig. 4). It has already been reported that Tn*5* insertion mutants in genes involved in core LPS synthesis or in O157 antigen synthesis, showed a reduced biofilm phenotype (34). An association between inactivation of LPS core synthesis genes and a decrease of biofilm formation was also observed in uropathogenic *E. coli* and *E. coli* K-12 (35). The biofilm negative mutants here have an altered LPS sugar composition and exhibit the deep rough pattern of LPS. A study conducted by Nakao and collaborators revealed that the *hldE* and *waaC* mutants involved in core OS biosynthesis exhibited a deep-rough LPS phenotype and an increase of biofilm formation (23).

### LPS of the Δ*pst* mutant are structurally different compared to the wild-type strain

The LPS structure of the Δ*pst* mutant is modified compared to its wild-type *E. coli* O157:H7. Mass spectrometry results confirmed that LPS of the Δ*pst* mutant are structurally different compared to EDL933 and culture conditions were shown to contribute to structural variability. The following major features were observed: 1) the absence of O-antigen in the mutant and the presence of an additional structure corresponding to 365.4u; 2) non acetylation of the core lateral GlcN is increased in M9 conditions for both strains; 3) PEA substitution of the lipid A phosphate groups is culture dependent for the wild-type strain and not culture dependent for the mutant; 4) lipid A palmitoylation is increased under biofilm conditions for both strains. Therefore, the Δ*pst* mutant produces rough-type LPS. Interestingly, the O157 antigen is absent upon activation of PagP, an outer membrane enzyme responsible of lipid A palmitoylation (36). Moreover, it was shown that biofilm-associated lipid A palmitoylation is a general feature of Enterobacteriaceae and this was associated to PagP activation in *E. coli* mature biofilm (25). Interruption of the O-antigen in the Δ*per* mutant of *E.coli* O157:H7 has previously been associated with increased autoagglutination (37) and increased adherence to Hela cells (38). This is in accordance with the autoagglutination phenotype of Δ*pst*. The addition of elements to core structures, blocking the addition of the O-chain has already been described in *Bordetella* species (39, 40).

Notably, the double mutant Δ*pst* Δ*phoB* that lacked the O-antigen LPS chain showed a low biofilm forming phenotype similar to the wild-type strain EDL933 but did not decrease in autoagglutination (Fig. 1B). Moreover, O-antigen was present in EDL933 grown in low Pi conditions. This suggests that the absence of O-antigen is independent of PhoB activation and could be associated with autoagglutination. The absence of the O-antigen might facilitate autoagglutination of the bacteria by exposing surface molecules involved in the enhanced phenomenon.

Confocal microscopy analyses of Δ*pst* mutant biofilm revealed that its biofilm matrix contained GlcNAc residues. But, these residues were absent in the biofilm matrices of all mutants with transposon inserted in LPS biosynthesis genes (Fig. 2 and Fig. 4). This suggests that detection of GlcNAc in the biofilm matrix was linked to LPS structure of the Δ*pst* mutant (Fig. 4). In *E. coli* O157:H7, the R3 core type LPS contains two molecules of GlcNAc (22): one linked to the heptose III from the inner core by *wabB* and the other linked to the galactose molecules from the outer core by an unknown protein (22). In addition, deletion of *pga* operon responsible for poly-N-acetylglucosamine production in mutant Δ*pst* (double mutant Δ*pst* Δ*pgaABCD*) did not influence the biofilm phenotype nor GlcNAc production (Fig. S5). This discards the contribution of poly-N-acetylglucosamine production to the GlcNAc composition of the Δ*pst* mutant biofilm matrix. Although non acetylation of the core lateral GlcN is increased in M9 conditions for both strains (Fig. 6), we propose that detection of GlcNAc in the biofilm matrix could be due to the absence of O157 molecules allowing the core LPS to be more exposed at the surface.

With the nutrient-limitation stress of phosphate limitations, activation of Pho and general stress regulons elicits specific and non-specific responses, respectively (41). Under Pi-starvation conditions the LPS biosynthesis, export genes, and O-antigen biosynthesis genes were differentially expressed in *E. coli* O157:H7 and in the Δ*pst* mutant. We demonstrated that PhoB activation was not related to the absence of O-antigen. In addition, several putative Pho boxes were found upstream of operons and genes that are involved in LPS biosynthesis and export (Table 2). To deal with the stress of phosphate starvation bacteria rely on different mechanisms to optimize the acquisition and bioavailability of phosphate and to maintain essential biochemical reactions. In this context, modifications of cell wall components such as LPS structures are part of the reorganization of the bacterial cell. Klein and collaborators showed that PhoB can regulate LPS heterogeneity (26, 42). In our study major structural variations were observed in LPS structures in the Δ*pst* mutant. In addition to the absence of O-antigen and the presence of an additional structure corresponding to 365.4u, the Δ*pst* mutant exhibited a constitutive PEA substitution of the lipid A phosphate groups (Fig. 7). We previously found that lipid A and fatty acid compositions are modified in the Δ*pst* mutant of avian pathogenic *E. coli* (43, 44).

### WaaH is a glycosyltransferase involved in biofilm formation

We also show here that the *waaH* (*yibD*) gene under PhoB control contributes to biofilm formation during phosphate starvation of the wild-type strain EDL933 and the Δ*pst* mutant. Its expression was strongly induced in both *E. coli* O157:H7 grown in phosphate-starvation conditions and in Δ*pst* mutant (Table 2). Interestingly, WaaH does not participate in the autoagglutination phenotype observed in Δ*pst* mutants. WaaH has been shown to be responsible for the addition of GlcUA on the inner core LPS of K-12 under phosphate-starvation conditions and under the control of PhoB. GlcUA modification was also observed in *E. coli* B, R2 and R4 core types and in *Salmonella* but not in *E. coli* O157:H7 R3 core (26). The addition of GlcUA on LPS core type was not detected by mass spectroscopy in LPS extracted from either EDL933 or from the Δ*pst* mutant. In *E. coli* O157:H7, the R3 core type, heptose III of its R3 core is decorated with GlcNAc by the plasmid-encoded gene *wabB* (Fig. 3). This precludes the addition of GlcUA on Heptose III (22, 26). Although the activity of WaaH in *E. coli* O157:H7 remains unknown, WaaH belongs to the glycosyltransferase 2 family and is conserved across the Proteobacteria phylum. Interestingly, WaaH shares 34% similarity with the glycosyltransferase PgaC, responsible for PGA elongation by adding GlcNAc molecules (26, 45). Thus, WaaH contributes to biofilm formation upon phosphate starvation and PhoB induction, its mechanism remains to be deciphered, but WaaH could be an interesting target for controlling biofilm formation.

## CONCLUSION

Various lines of evidence suggest that the Pho regulon and key factors involved in nutritional and other stress responses are interrelated. To survive adverse conditions that may play a role in its transmission, persistence, and virulence *E. coli* O157:H7 has developed strategies such as biofilm formation involving the Pho regulon.

## MATERIALS AND METHODS

### Bacterial strains

The effect of the Pho regulon on EHEC O157:H7 biofilm formation was investigated using strain EDL933 and its derivative mutants (listed in Table 1). The EDL933 Δ*pst* and Δ*phoB* mutants were previously constructed (46). The Δ*waaH* single mutant and Δ*pst* Δ*phoB* or Δ*pst* Δ*waaH* double mutants were generated by allelic exchange of *phoB* or *waaH* genes using a suicide vector in EDL933 wild-type strain or in EDL933 Δ*pst* mutant as previously described (47) with some modifications. Using the primers listed in Table S1, a tetracycline resistance cassette from pACYC184 was amplified and flanked by PCR with the 500 bp sequence adjacent to the *phoB* or *waaH*. Tetracycline was used to select the Δ*waaH* mutant, and Δ*pst* Δ*waaH,* or Δ*pst* Δ*phoB* double mutants. The deletion of *pgaABCD* operon was generated in EDL933 Δ*pst* using the lambda Red recombinase system (48) by recombining a chloramphenicol cassette from the pKD3 in the *pga* locus of EDL933 Δ*pst*. The deletion of *phoB*, Δ*pgaABCD*, and *waaH* were confirmed by PCR and sequencing. Complementation of *waaH* was performed using the pTrc99a expression plasmid. The *waaH* ORF was amplified from the genome of EDL933 using the primers containing BamHI and SacI site of digestion as listed in Table S1. *waaH* was then inserted into pTRC99a downstream of the isopropyl-β-D-thiogalactopyranoside (IPTG) inducible pTrc promoter. As complementation was observed without induction by IPTG, IPTG was not added to the media. LB cultures were made from one isolated colony at 37°C on LB agar containing antibiotics when required and incubated for 24 h at 37°C. Antibiotics were used at the following concentrations: 10 μg/ml tetracycline (Tc); 50 μg/ml ampicillin (Amp); 25 μg/ml chloramphenicol (Cm); and 50 μg/ml kanamycin (Km).

### Static biofilm formation assay

The biofilm formation was investigated as described previously (49) with modifications. Overnight cultures at 37°C in LB media were diluted (1:100) in 5 ml of morpholine propanesulfonic acid (MOPS) minimal medium (Teknova) containing 0.2% (wt/vol) glycerol, thiamine (0.1 μg/mL), and 1.32 mM (MOPS high Pi) KH_2_PO_4_ or in M9 medium with 0.2% (wt/vol) glycerol and minerals (1.16 mM MgSO_4_, 2 µM FeCl_3_, 8 µM CaCl_2_, and 16 µM MnCl_2_) and incubated for 24 h at 37°C with antibiotics when required. Effect of high and low phosphate concentration on EDL933 biofilm formation was studied using MOPS culture dilution (1:100) either in MOPS with 1.32 mM (MOPS high Pi) KH_2_PO_4_ or 1 µM (MOPS low Pi) KH_2_PO_4_. Biofilm formation of EDL933 and its derivative mutants was studied using M9 medium dilutions (1:100) with 0.2% (wt/vol) glycerol and minerals (1.16 mM MgSO_4_, 2 µM FeCl_3_, 8 µM CaCl_2_, and 16 µM MnCl_2_) without use of antibiotics. These dilutions were inoculated in triplicate into microtiter plates (Costar® 3370; Corning, NY, USA). After 24 h of incubation at 30°C, unattached cells were removed by washing three times with distilled water. Plates were dried at 37°C for 15 min and biofilms were stained with 0.1% (wt/vol) crystal violet for 2 min. The crystal violet solution was removed and biofilms were washed three times with distilled water and then dried at 37°C for 15 min. The stain was released with 150 μl of 80% (vol/vol) ethanol and 20% (vol/vol) acetone. The biofilms were quantified by measuring the absorbance at 595 nm with a microplate reader (Powerwave; BioTek Instruments, Winooski, VT, USA). Moreover, while biofilms were stained with crystal violet and measured with the A_595_, growth of the planktonic part of the culture (unattached cells) was tracked with OD_600_. Relative biofilm formation corresponds to A_595_/OD_600_ (50). One-way ANOVA with Dunnett’s multiple-comparison post hoc tests were performed to calculate *p*-values.

### Autoagglutination assays

Autoagglutination was determined using an assay described by Bansal et al (51) with modifications. EDL933 and its isogenic mutant were cultivated in LB media at 37°C and diluted 1/100 in M9 and incubated at 30°C for 24 h without agitation. After incubation, 100 µl of the upper phase was taken 1 cm below the surface and the OD_600_ was measured (OD_Agg_). Cultures were then mixed by vortex for 10 s and OD_600_ was measured again (OD_tot_). The percentage of autoagglutination was calculated by using this formula: [(OD_tot_-OD_Agg_) / OD_tot_] x 100. One-way ANOVA with Dunnett’s multiple-comparison post hoc test was performed to calculate *p*-values.

### Confocal laser scanning microscopy

Stained biofilms were visualized by confocal laser scanning microscopy at 40X (CLSM; FV1000 IX81; Olympus, Markham, ON, Canada) as previously described (49).

### Screening of Tn*10* Δ*pst* mutant library

Km^R^ Cm^R^ independent mutants were screened for the loss of autoagglutination. This assay is based on the ability of the Δ*pst* mutant to autoagglutinate and to form a larger ring at the bottom of a 96-well round microtiter plate compared to wild type (Fig. 1 and Fig. S3). As autoagglutination phenotypes were similar in LB and M9, LB was chosen to facilitate manipulations. Each plate was filled with 150 µl of LB broth and filled with one colony of Δ*pst* mutant containing Tn*10* insertions. In each plate, EDL933 and Δ*pst* mutant were used as negative and positive controls of agglutination phenotype respectively, and wells containing LB only was used as a blank. After 24 h of incubation at 30°C with agitation, the ring sizes were eyes checked and mutants that loose the ability to form large rings were evaluated two more times for autoagglutination. Those mutants were evaluated for their biofilm formation in M9 broth at 30°C for 24 h, as described above. Mutants that lost the ability to form biofilm similar to the Δ*pst* mutant were stored at −80°C for further analysis. In order to avoid mutants with growth defects, OD_600_ was measured for 24h of incubation at 30°C in M9 broth (Fig. S4).

### Identification of mini-Tn*10* insertion sites

Mini-Tn*10* insertion sites of Δ*pst* biofilm negative mutants were identified by DNA sequencing. To extract DNA, each transposon mutant was cultivated separately in LB media and 1 ml of 7 to 10 clones belonging to the same phenotype group were pooled. Genomic DNA was extracted with Qiagen DNeasy blood and tissue kits and pooled at an equimolar concentration in order to prepare a total of three sequencing libraries using KAPA Hyper Prep kit. The libraries were then sequenced using an Illumina MiSeq apparatus at the Plateforme d’Analyses Génomiques of the Institut de Biologie Intégrative et des Systèmes (IBIS, Université Laval). The resulting sequencing reads were mapped on the extremities of the sequence of the Tn*10* transposon using bwa version 0.7.12-r1039 (52). The mapped reads were subsequently converted to fastq format with SAMtools version 0.1.19-44428cd (52) and the Tn*10* transposon sequence was filtered from the reads using cutadapt version 1.10 (53). The resulting filtered reads were mapped using bwa on the genomic sequence of *E. coli* O157:H7 EDL933 (GenBank: AE005174.2). The features about where the reads mapped were found using featureCounts (54), which is included in the package subread version 1.5.0-p3. The presence of the transposon in intergenic regions was investigated using Artemis version 16.0.0 (55). Finally, categories of orthologous groups were analyzed with eggNOG-mapper.

### Microarray experiment

EDL933 and its isogenic Δ*pst* mutant were grown in high Pi MOPS 0.2% (wt/vol) glucose condition until the OD600 nm reached 0.6 (56). Samples equal to 5 × 10^8^ CFU were taken from this mid-log phase and processed for transcriptome analysis. One µg of the fragmented and biotinylated cDNAs, were hybridized onto Affymetrix GeneChip^®^ *E. coli*. The data were processed using the FlexArray^®^ software; Robust Multi-array Average (RMA) normalization. The levels of transcription obtained from 3 biological replicates of each experimental condition were compared using the EB (Wright & Simon) algorithm. The comparison was conducted between the Δ*pst* mutant and the wild-type strain with the differential expression conditions corresponding to a 2-fold change (FC) cut-off and *p-*value < 0.05. Validation of microarray results was achieved by comparing the expression of eleven genes by qRT-PCR. The 16S rRNA gene, *tus*, was included for normalization within samples.

### *In silico* evaluation of Pho boxes

The presence of putative Pho boxes upstream of biofilm-related genes was evaluated with a previously constructed Pho box matrix (21) and was based on 12 known Pho boxes with O157:H7 EDL933 sequences (57). Using MEME suite website with the FIMO tools version 4.12.0 (http://meme-suite.org/tools/fimo), nucleotide sequences of LPS promoting regions were scanned to find putative motif sequences.

### LPS extraction

To extract LPS from planktonic and biofilm cells of EDL933 and its isogenic mutants, each strain was diluted (1/100) and grown in 6 Petri dishes containing 20 ml of M9 media for 24 h at 30°C. Planktonic cells (20 ml) were kept for further analyses. The 6 biofilms were washed with 20 ml of PBS. PBS (1 ml) was added in each plate and biofilms were scratched from the surface with a cell scratcher. Biofilm and planktonic cells were pelleted by centrifugation at 4000 rpm for 15 min and suspended to an OD_600nm_ of 0.6. Then, 1.5 ml of suspension was pelleted in a 2 ml Eppendorf tube at 13.000 rpm for 3 min. The supernatant was discarded and the pellet was stored at −20°C before LPS extraction. LPS was extracted as previously described (58). Briefly, a defrosted pellet was suspended in 50 µl of SDS-buffer (4% β-mercaptoethanol (BME), 4% SDS, and 10% glycerol in 0.1 M Tris-HCl, pH 6.8) and boiled for 10 min. After cooling down, 5 µl of 10 mg/ml solution of DNase I in DNase buffer (150 mM NaCl and 1 mM CaCl2) and 5 µl of 10 mg/ml RNase A in sterile water was added to samples and incubated at 37°C for 30 min. After adding 5 µl of 20 mg/ml proteinase K in proteinase K buffer (50 mM Tris-HCl and 1 mM CaCl_2_), samples were incubated at 60°C for 1 hour. Each sample was treated with phenol by adding 50 µl of ice-cold Tris-saturated phenol and by incubating at 65°C for 15 min. Phenol treated samples were mixed by vortex every 5 min. Samples were cooled to room temperature and 500 µl of diethyl ether were added and mixed by vortex for 10 s. Samples were centrifuged at 20,600 x g for 10 m and the bottom phase was carefully removed. Phenol treatments were repeated at least three times. Alternatively, for MALDI-MS analyses, LPS were extracted as described in the MALDI-MS section.

### SDS-PAGE and Western blot analysis

Prior to gel electrophoresis, 100 µl of 2X SDS-buffer containing bromophenol blue 0.002% (wt/vol) were added to the extracted sample. 10 µl of LPS extracted sample were separated by electrophoresis on 15% SDS-PAGE at 100 volts for 18 min (upper gel) and then 200 volts for 60 min (lower gel). LPS samples were visualized by using the gel silver staining procedure according to the instructions of the manufacturer (Biorad). Briefly, the gel was covered with a fixative solution (methanol 40% and acid acetic 10% (vol/vol)). After overnight incubation, the gel was covered with 10% (vol/vol) oxidizer for 5 min and then washed 6 to 7 times with water for 15 min. The gels were then exposed to 10% (vol/vol) silver reagent for 20 min. After a 30 s wash in water, gels were developed in 3.2% Developer solution (wt/vol). LPS were also visualized by Western blot. After electrophoresis, polyacrylamide gel was stabilized by covering it with 1X transfer buffer (25 mM Tris-base pH 8.2, 192 mM glycine and 20% (vol/vol) methanol) for 30 min. LPS were then transferred to nitrocellulose membranes (Biorad) at 14 volts. After overnight transfer, the nitrocellulose membrane was blocked in T-TBS with 1% skimmed milk for 1 h. The membrane was washed 3 times with T-TBS and incubated with 1/1000 solution of primary O157:H7 monoclonal antibody (LifeSpan BioSciences) for 1 h 40 min. The membrane was washed three times with T-TBS for 3 x 5 min and incubated for 1 h with 1/1000 solution of horseradish peroxidase (HRP)-conjugated goat anti-mouse IgG antibody (1:1000; Biorad) and washed 3 times with T-TBS and 1 time with TBS. The antigen-antibody reaction was analyzed by using Opti-4CN detection kit according to the instructions of the manufacturer (Biorad).

### Mass spectrometry analyses

LPS were extracted from different growth cultures of the wild-type EDL933 strain and its Δ*pst* mutant. The LPS-BioSciences proprietary LPS extraction method (59) was applied to 10 mg quantities of lyophilized bacterial pellets issued from the six bacterial cultures under study, i.e. EDL933 and Δ*pst* LB, M9 planktonic and M9 biofilm cultures. The non-discrimination of molecular species by the extraction procedure were verified by comparison of SDS PAGE profiles of the extracted LPS with those obtained from corresponding non-fractioned bacterial lysates (60).

Polysaccharide (PS) and lipid A moieties of each LPS were analyzed separately by MALDI MS. Prior to the analyses, crude LPS samples were cleaved under mild acid hydrolytic conditions established in (61). Hydrolysates were lyophilized, and lipid A and PS moieties were extracted subsequently, first with a chloroform-methanol-water mixture for the lipids A, and then with water for the corresponding PS.

Each lipid A and PS samples were mixed on the MALDI target with appropriate matrix solutions and let dry. Di-hydroxy-benzoic acid (DHB) was used as a matrix. For lipid A analysis, DHB was dissolved at 10 µg/µl in 0.1 M citric acid in a chloroform-methanol water mixture and for PS analysis in 0.1 M citric acid in water. Different analyte/matrix ratios were tested.

MALDI MS analyses were performed in linear the negative-ion mode on a Shimadzu AXIMA Performance mass-spectrometer. Negatively charged [M-H]^−^ molecular ions were desorbed by a 337 nm N2 laser pulses and analyzed in the linear mode. 1000 laser shots were accumulated for each sample (61).

### Data availability

Microarray data have been deposited in the Gene Expression Omnibus database https://www.ncbi.nlm.nih.gov/geo/query/acc.cgi?acc=GSE125488 under accession number GSE125488.

## ACKNOWLEDGMENTS

We thank Dr. Yannick D.N. Tremblay for sharing his expertise and technical assistance and Frederic Berthiaume for his help with confocal laser scanning microscopy assays. We are grateful to Judith Kashul for editing the manuscript. This work was supported by grants from the Natural Sciences and Engineering Research Council of Canada (RGPIN SD-25120-09) to Josée Harel and Fonds de la recherche du Québec en nature et technologies to Josée Harel and Mario Jacques (FRQNT PT165375) and a scholarship to Philippe Vogeleer from FRQNT Québec Wallonie program (FRQNT Regroupements stratégiques 111946).

